# Multiple Myeloma DREAM Challenge Reveals Epigenetic Regulator *PHF19* As Marker of Aggressive Disease

**DOI:** 10.1101/737122

**Authors:** Mike J Mason, Carolina Schinke, Christine L P Eng, Fadi Towfic, Fred Gruber, Andrew Dervan, Brian S White, Aditya Pratapa, Yuanfang Guan, Hongjie Chen, Yi Cui, Bailiang Li, Thomas Yu, Elias Chaibub Neto, Konstantinos Mavrommatis, Maria Ortiz, Valeriy Lyzogubov, Kamlesh Bisht, Hongyue Y Dai, Frank Schmitz, Erin Flynt, Dan Rozelle, Samuel A Danziger, Alexander Ratushny, Multiple Myeloma DREAM Consortium, William S Dalton, Hartmut Goldschmidt, Herve Avet-Loiseau, Mehmet Samur, Boris Hayete, Pieter Sonneveld, Kenneth H Shain, Nikhil Munshi, Daniel Auclair, Dirk Hose, Gareth Morgan, Matthew Trotter, Douglas Bassett, Jonathan Goke, Brian A Walker, Anjan Thakurta, Justin Guinney

## Abstract

While the past decade has seen meaningful improvements in clinical outcomes for multiple myeloma patients, a subset of patients do not benefit from current therapeutics for unclear reasons. Many gene expression-based models of risk have been developed, but each model uses a different combination of genes and often involve assaying many genes making them difficult to implement. We organized the Multiple Myeloma DREAM Challenge, a crowdsourced effort to develop models of rapid progression in newly diagnosed myeloma patients and to benchmark these against previously published models. This effort lead to more robust predictors and found that incorporating specific demographic and clinical features improved gene expression-based models of high risk. Furthermore, post challenge analysis identified a novel expression-based risk marker and histone modifier, *PHF19*, which featured prominently in several independent models. Lastly, we show that a simple four feature predictor composed of age, International Staging System stage (ISS), and expression of *PHF19* and *MMSET* performs similarly to more complex models with many more gene expression features included.

**Key points:**

- Most comprehensive and unbiased assessment of prognostic biomarkers in MM resulting in a robust and parsimonious model.
- Identification of *PHF19* as the expression based biomarker most strongly associated with rapid progression in MM patients.

## Introduction

Multiple myeloma (MM) is a hematological malignancy of terminally differentiated plasma cells (PCs) that reside within the bone marrow^1^. It arises as a result of complex chromosomal translocations or aneuploidy with approximately 25,000-30,000 patients diagnosed annually in the United States^2,3^. The disease’s clinical course depends on a complex interplay of molecular characteristics of the PCs and patient socio-demographic factors. While progress has been made with novel treatments extending the time to disease progression (and overall survival) for the majority of patients, a subset of 15%-20% of newly diagnosed MM patients are characterized by an aggressive disease course with rapid disease progression and poor overall survival regardless of initial treatment^4–6^. Accurately predicting which newly diagnosed patients are at high-risk is critical to designing studies that will lead to a better understanding of myeloma progression and facilitate the discovery of novel therapies that meet the needs of these patients.

To date most MM risk models use patient demographic data, clinical laboratory results and cytogenetic assays to predict clinical outcome. High risk defining cytogenetic alterations typically include deletion of 17p and gain of 1q as well as t(14;16), t(14;20), and most commonly t(4;14), which leads to juxtaposition of *MMSET* with the immunoglobulin heavy chain locus enhancer, resulting in overexpression of the *MMSET* oncogene^4^. While cytogenetic assays, in particular fluorescence *in situ* hybridization (FISH), are widely available, their risk prediction is sub-optimal and recently developed classifiers have used gene expression data to more accurately predict risk^7–9^. To investigate possible improvements to models of newly diagnosed myeloma progression, we organized the crowd sourced Multiple Myeloma DREAM Challenge, focusing on predicting high-risk, defined as disease progression or death prior to 18 months from diagnosis. This benchmarking effort combined eight datasets, four which provided participants with clinical, cytogenetic, demographic and gene expression data to facilitate model development, and four hidden, independent data sets (N = 823 unique patient samples) for unbiased validation. Over 800 people participated in this Challenge, resulting in the submission of 171 predictive algorithms for objective evaluation. Several models submitted to the Challenge demonstrated improved accuracy over existing state-of-the-art, published models.

Analysis of top performing methods identified high expression of *PHF19*, a histone methyltransferase, as the gene with the strongest association with myeloma progression, with greater predictive power than the expression level of the known high risk gene *MMSET*. We developed a four parameter model using age, ISS, and *PHF19* and *MMSET* expression that performs as well as more complex models having many more gene features. The parsimony of this model should facilitate its translation to the clinic. Significantly, we showed that knock down of *PHF19* shifts myeloma cell lines into a less proliferative state. To our knowledge, this is the first DREAM Challenge to both nominate and experimentally validate a candidate biomarker and, as such, demonstrates the biological and clinical impact of crowdsourced efforts.

## Methods

### Datasets

The Challenge includes five microarray and three RNA-seq expression datasets, annotated with clinical characteristics such as gender, age, ISS, and cytogenetics (Table 1)^9–14^. In all datasets, the expression assay was performed on CD138+ PCs isolated from bone marrow aspirates or blood of newly diagnosed patients. Data were split into training and validation datasets (Table 1).

**Table 1:**
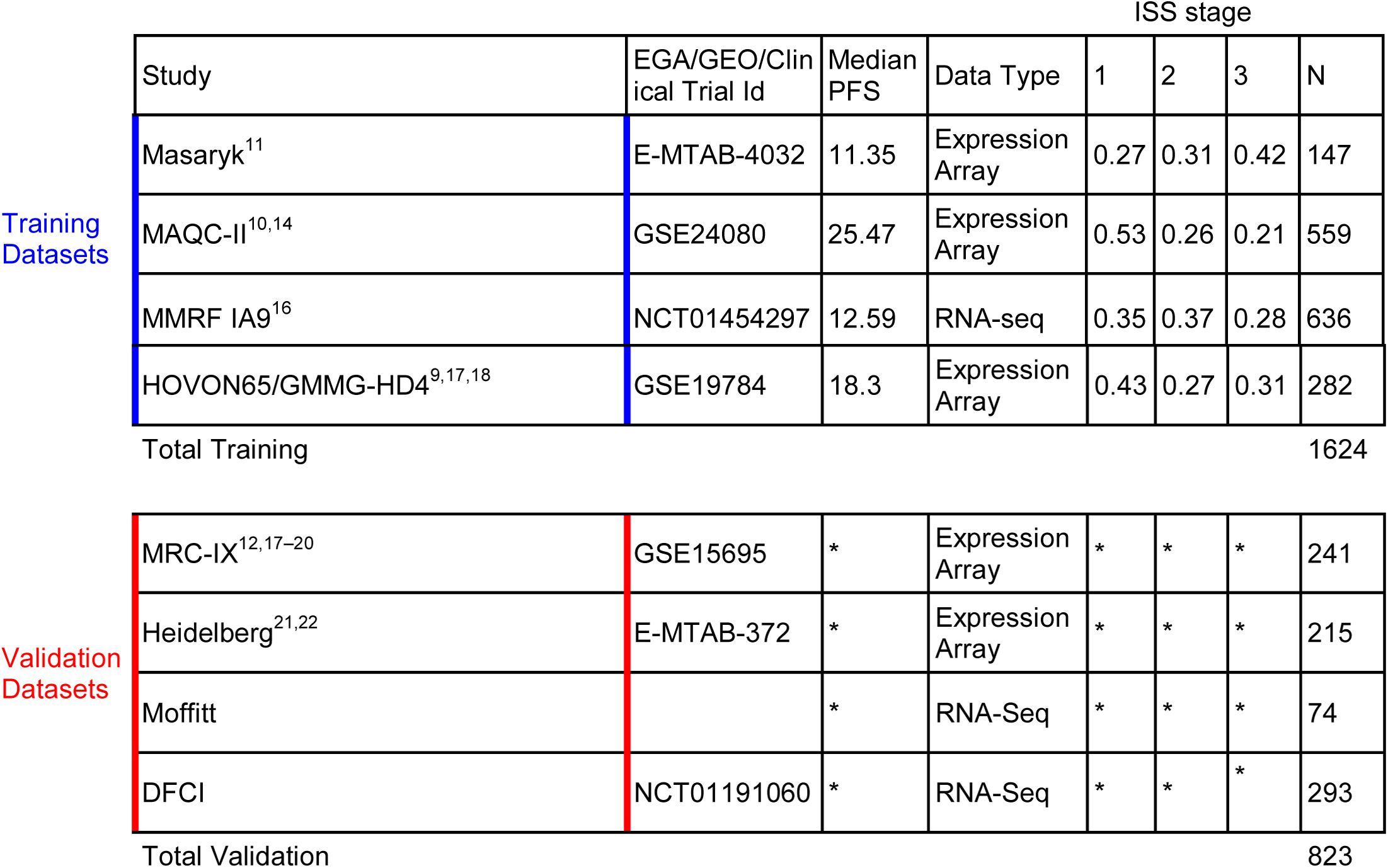
Data set descriptions. *: clinical data withheld per data provider request.

Three institutes provided RNA-sequencing data. The Multiple Myeloma Research Foundation (MMRF) provided an additional training dataset from its publicly available CoMMpass study (release IA9). Collaborating with the Myeloma Genome Project / Dana Farber Cancer Institute (MGP-DFCI) access was provided to data from their clinical trial where patients were randomized into a standard treatment arm and an aggressive treatment arm that included autologous stem cell transplant (ASCT) and high dose therapy^15^. An additional dataset from the Oncology Research Information Exchange Network (ORIEN) was made available through a collaboration with Moffitt Cancer Center and M2Gen (See supplement for more details on datasets).

### Assessing Model Performance

To identify top performing teams we employed two metrics to assess the accuracy of submitted models within a given validation cohort: the integrated area under the curve (iAUC) and balanced accuracy curve (BAC). While the AUC is a widely accepted metric of prediction accuracy, it is sensitive to the specific time threshold used to differentiate high and low patient risk. The myeloma research community has not yet reached a consensus on the time point that best separates patients into risk groups, though there is a general agreement that it lies between 1 and 2 years for progression free survival (PFS). We therefore chose the more robust iAUC for the primary metric, with a sliding PFS threshold between 12 and 24 months at weekly increments. iAUCs computed in each validation cohort were combined into a weighted average (wiAUC) with each cohort iAUC weighted by the square of the number of high-risk patients.

Using the wiAUC we computed the Bayes factor, *K*, to identify statistically tied top performing predictors (see supplemental Methods). Predictors with *K*_*p*_< *3* are considered tied with the top scoring model and the weighted BAC (wBAC) was used as a tie-breaking metric in order to determine the final top performing model, with weighting by the square of the number of high risk patients in each dataset (see supplemental text).

### *In vitro* studies For Functional assessment of *PHF19*

Studies to determine the functional importance of PHF19 were performed using standard assays and are described in the supplemental text. In brief, tetracycline inducible short hairpin RNA (TRIPZ shRNA) was used to knockdown *PHF19* expression in two MM cell lines (JJN3 and ARP1). A non-silencing scrambled TRIPZ shRNA was used as a control. *PHF19* knockdown (KD) after doxycycline induction was measured by quantitative real time polymerase chain reaction (qRT-PCR) and western blotting. Cell viability (Cell Titer Blue, Promega), cell cycle (Vybrant DyeCycle Stain, Thermo Scientific) and apoptosis (Annexin V, Biolegend) were assessed and differences between the *PHF19* KD cells and control group were analyzed.

## Results

### Top Challenge Models Outperform Baseline and Published Myeloma Predictors

To develop and assess prognostic models of high-risk in MM, we assembled eight data sets totaling 2447 patients annotated with overall survival (OS) and progression-free survival (PFS; Table 1). We asked participants to predict whether a patient was high-risk as defined as disease progression or death prior between 12-24 months since diagnosis (see Methods). Participants developed prognostic models using clinical features (e.g., age, sex, International Staging System (ISS), and cytogenetic features) and gene expression utilizing four training datasets. Challenge participants submitted models to be evaluated against four validation cohorts sequestered in the cloud (Figure 1, Table 1, see supplement text). Model predictions were benchmarked against each other and comparator models (Table 2) using weighted-integrated AUC (wiAUC), with statistical ties resolved using weighted balanced accuracy (wBAC) (see methods and supplemental text).

**Figure 1.**
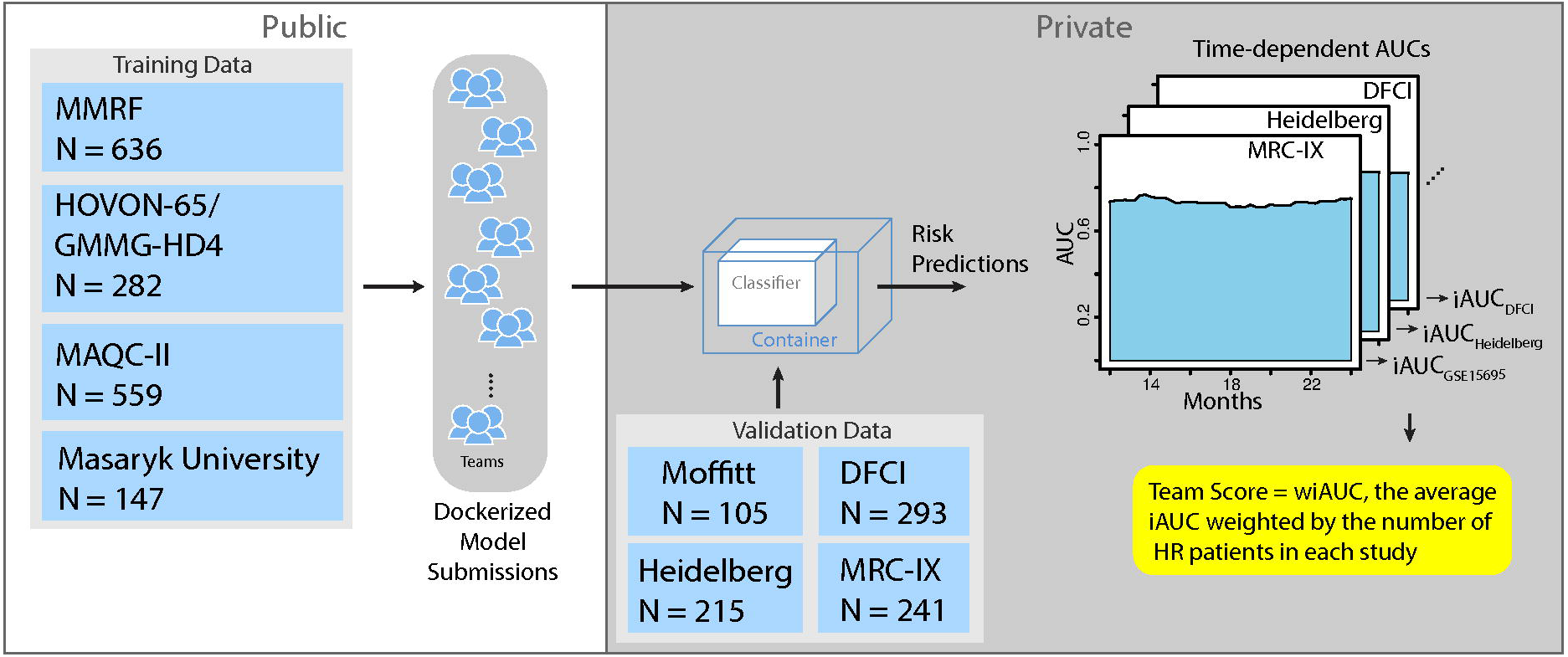
Challenge Model Submission Architecture: training datasets are fully available to challenge participants (left), while validation datasets are sequestered in the cloud (right). Containerized models are submitted to cloud, ran on training datasets and risk predictions are scored.

**Table 2:** Comparator modes.

| Model | Reference | Features |
| --- | --- | --- |
| Baseline | Baseline | Age and/or ISS |
| UAMS-70 | Shaughnessy *et al* 2007<sup>7</sup> | Gene expression signature composed of 70 genes |
| EMC-92 | Kuiper *et al* 2012<sup>23</sup> | Gene expression signature composed of 92 genes |
| UAMS-70 extended | This manuscript | UAMS-70 with age and/or ISS |
| EMC-92 extended | This manuscript | EMC-92 with age and/or ISS |

Of 42 finalized models submitted to the Challenge, 11 exceeded the performance of the age plus ISS baseline model (Figure 2, wiAUC=0.6207). The top-performing predictor, developed by researchers at the Genome Institute of Singapore (GIS), outperformed all Challenge participant models (wiAUC = 0.6721) as well as the published comparator models UAMS-70 (wiAUC=0.6414) and EMC-92 (wiAUC=0.6042, Figure 2).

**Figure 2.**
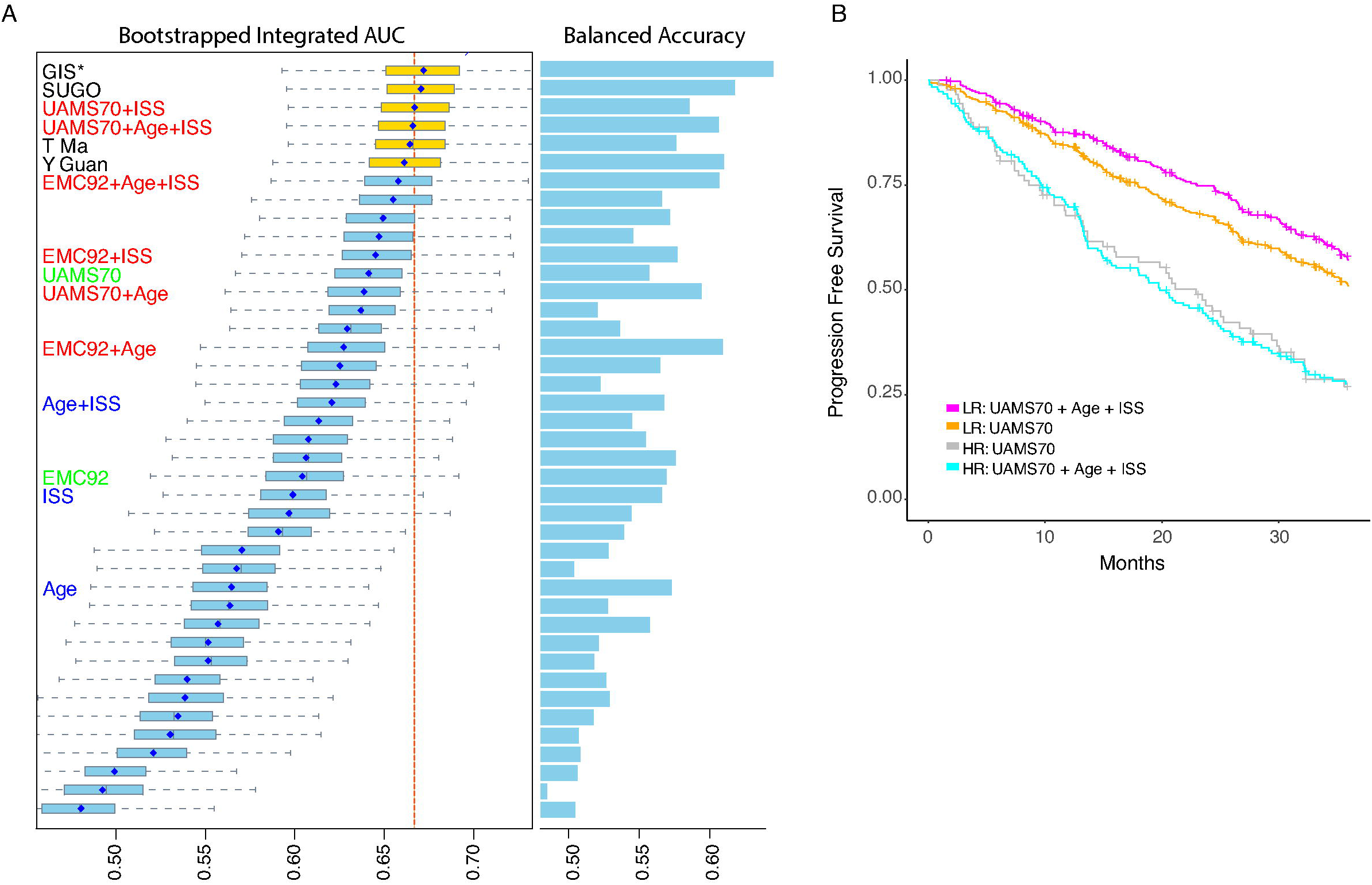
Challenge Performance: A) Box plots show distributions of bootstrapped model performances for each team. Comparator models are shown with text marked in blue for baseline models, green for published models and red for published models extended to include clinical features. The dashed red line indicates the median of the best performing comparator model. Barplots to the right show the tie-breaking metric, wBAC, for each model. Amongst statistically tied models, GIS has the highest wBAC and was declared the top-performer. Asterisk indicates internal collaborator’s comparator model. B) Kaplan-Meier curve of UAMS-70 comparator model with and without age and ISS added.

### Combining Clinical Features and Gene Expression Features Improves Myeloma Risk Prediction Accuracy

After the Challenge submission period ended, Challenge organizers and top performing teams assessed which features had the largest impact on model performance. ISS was the most important model feature in GIS’s top-performing model as measured by the mean decrease in Gini coefficient (see methods). A DNA repair signature previously associated with poor prognosis^24^ was the second most important feature, while age ranked third (supplemental Figure 1).

To assess whether age and/or ISS explained the difference in model performance between the top-performing model and published comparator models, we extended the UAMS-70^7^ and EMC-92^23^ models to include age and/or ISS and assessed their performance (Figure 2; see Supplemental Methods). The addition of these clinical features improved performance of both published models: EMC-92 wiAUC= from 0.6042 versus EMC-92+age+ISS wiAUC=0.658 and UAMS-70 wiAUC=0.6414 versus UAMS-70+ISS wiAUC=0.6667 as has been suggested previously^13,25^. Adding ISS to the UAMS-70 model improved its accuracy such that it was tied with the top-performing model (i.e., its Bayes factor *K* < 3; Figure 2a-b).

### Top-Performing Challenge Methods Identify *PHF19* as a Novel Myeloma High Risk Biomarker

The top-performing model implemented a wisdom of the crowd approach, utilizing clinical features and published myeloma signatures that summarize the expression of gene sets. The second-place SUGO model instead included individual genes as features, utilizing a univariate-based feature selection approach to identify genes to include in their model. In each of the four training datasets the SUGO team computed each gene’s effect size, *z*, via the concordance index between overall survival and the gene’s expression. These effect sizes were combined across training sets using Stouffer’s method^26^ without weighting to yield a single meta-*z* per gene. The meta concordance index was calculated using two expression normalization procedures, with one nominating *PHF19* as the most important gene and the second identifying *CDKN3* (See Supplement Methods).

We replicated this analysis in both the training and validation expression datasets using PFS in place of OS to increase statistical power. We also weighted studies according to their number of high risk patients when applying Stouffer’s method. This univariate analysis also found previously described myeloma risk genes *MMSET* and *CKS1B* with large values in the tail of the meta-*z* distribution (Figure 3a), validating this approach. The meta-*z* values of these genes were surpassed by *PHF19*, the top-ranked gene regardless of normalization procedure (not shown) in both the training and validation datasets (Figure 3a).

**Figure 3.**
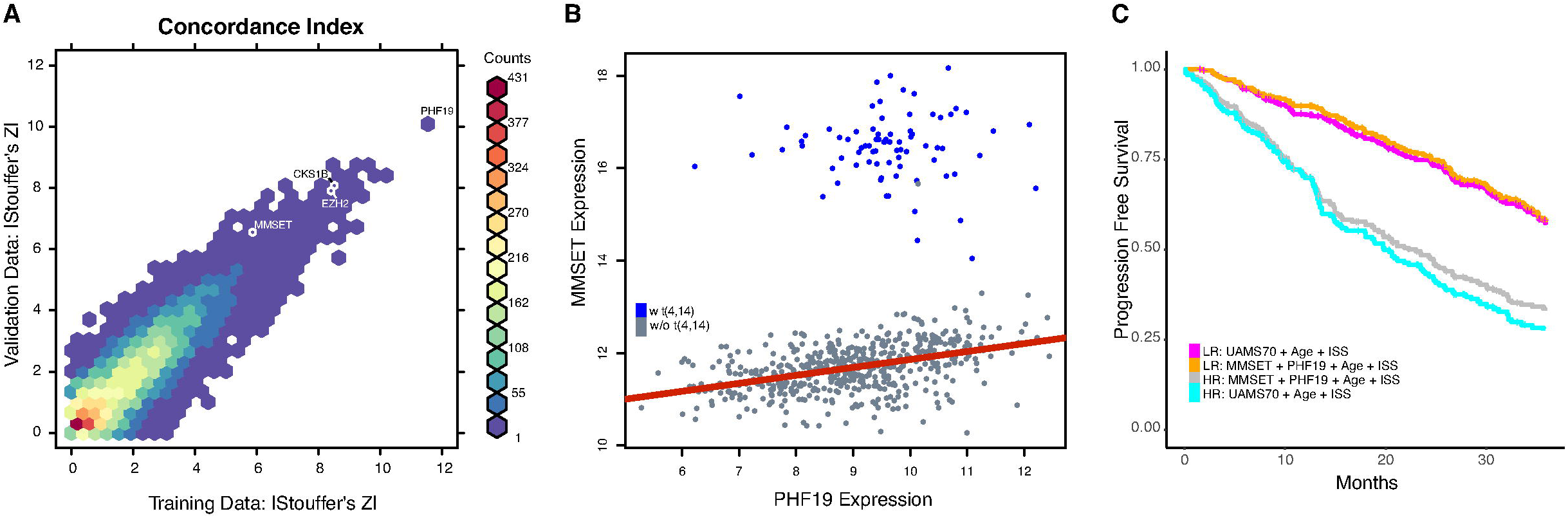
*PHF19* compared to other Myeloma classifiers and Features: A) Two dimensional histogram of PFS concordance index-based univariate effect sizes (z) in training and validation cohorts where colors represent the number of genes in a given hexagonal bin. *PHF19* and well-known myeloma genes noted. B) *PHF19* and *MMSET* expression in relation to t(4;14). C) A simple four feature model performs as well as UAMS-70 combined with age and ISS.

Given that *PHF19’s* association with progression was replicated in the validation datasets, we next asked whether it could improve the performance of the GIS model which did not include single genes. The GIS model uses penalized logistic regression over features ranked and selected based on mean decrease in Gini coefficient calculated via a random forest (see Supplemental Methods). We added *PHF19* expression into this feature selection process and found that it ranked higher than all other features, except ISS and a previously-published DNA repair signature^24^(supplemental Figure 2).

### Incorporating PHF19 and MMSET expression with Age and ISS Identifies a Simple Model of High Risk MM

Given that both *PHF19* and *MMSET* are histone modifiers playing a role in H3K36 methylation we checked whether their expression is correlated (Figure 3b). Expression of *MMSET* and *PHF19* do not appear to have an association. As has previously been shown, *MMSET* expression is clearly affected by the t(4;14) translocation with the immunoglobulin enhancer driving high *MMSET* expression, but expression of *PHF19* is not correlated. However, subsetting by t(4;14) reveals a modest linear relationship between *MMSET* and *PHF19* expression in samples lacking the translocation (*r = 0*.*423*) while there is no such relationship in the t(4;14) positive samples (*r = -0*.*067*).

Given the impact of *PHF19* on model performance and *MMSET*’s status as a reported myeloma risk marker, we checked to see if a model composed of age, ISS, *PHF19* and *MMSET* could perform as well as one using the features of the top-performing extended comparator model (UAMS-70 plus age plus ISS). We constructed a Cox proportional-hazards model of the two feature sets and found that the four parameter model (wiAUC=0.693) out-performed the UAMS-70-based model in the validation cohort (wiAUC=0.667; Figure 3c) placing it on par with the winning algorithm.

### Knockdown of *PHF19* Leads to Decreased Proliferation through Cell Cycle Arrest in Multiple Myeloma Cell Lines

To determine whether *PHF19* is functionally important for the malignant growth of MM cells, we used lentiviral-expressed shRNA directed against *PHF19*. We transduced JJN3 and ARP1 MM cell lines with a shRNA targeting *PHF19* or a scrambled control shRNA and selected out transduced cells. shRNA induced cells showed knockdown (KD) of >90% *PHF19* RNA and protein relative to the control after 72 hrs and 168 hrs for the JJN3 and ARP1 cell lines, respectively (Figure 4a and 4b). KD of *PHF19* led to significant inhibition of proliferation in the JJN3 and ARP1 MM cell lines compared to scrambled shRNA control (Figure 4c-d) confirming the recent finding of PHF19’s effect on proliferation in MM cell lines^27^. To identify the mechanism of growth inhibition, we performed cell cycle analysis and observed a significant arrest of MM cells in the G0/G1 stage with *PHF19* KD compared to the scrambled control shRNA (Supplemental Figure 3a). This was seen consistently in both cell lines examined (Supplemental Figure 3b). We further investigated the effect of *PHF19* KD on apoptosis and necrosis, but did not find significant differences at the examined time points (Supplemental Figure 3c-e). These results demonstrate that *PHF19* is functionally relevant in MM and that reduction of *PHF19* leads to a decrease in cell proliferation via cell cycle arrest.

**Figure 4.**
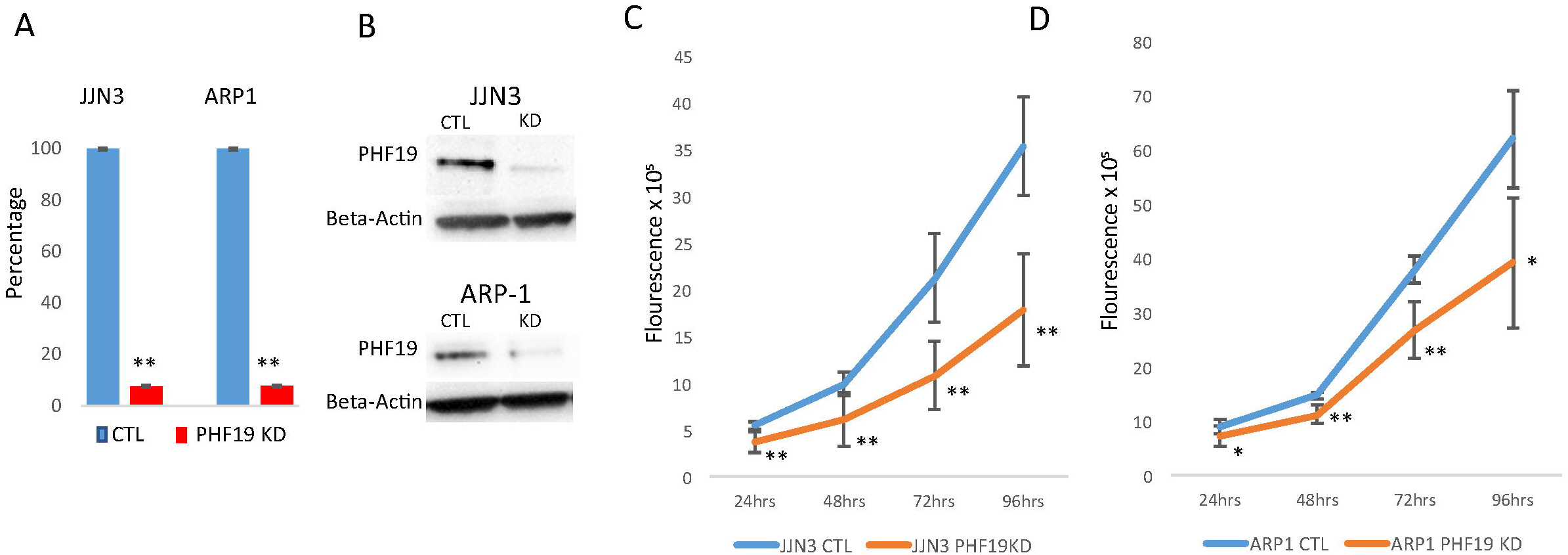
PHF19 knock down leads to decreased cell: Knockdown of PHF19 was performed in the JJN3 and ARP1 MM cell lines using inducible shRNA. A) PHF19 knockdown, relative to scrambled shRNA control, was confirmed using qRT-PCR and B) western blotting. C-D) Cell proliferation was significantly decreased in MM cells with PHF19 knockdown compared to scrambled control for JJN3 and ARP1 cell lines.

## Discussion

In the course of the crowd sourced Multiple Myeloma DREAM Challenge we benchmarked 171 prediction models and found that the accuracy of gene expression-based models benefited from the addition of clinical data, specifically: age and ISS improved AUC-based metrics by approximately 6% while an indicator of whether a patient received an ASCT improved the metric by roughly 5%. As such, expression-based patient stratification efforts should incorporate age and ISS, and possibly include an ASCT indicator for any *post hoc* analysis.

Additionally, we show for the first time that expression of *PHF19* is a stronger predictor of MM progression than the expression level of the high risk marker *MMSET*, which is particularly overexpressed in patients with the high risk translocation t(4;14). This strong association was likely missed in earlier studies given that *PHF19* expression is not associated with any cytogenetic feature while several therapeutic advances over the last 20 years have made it difficult to model outcome across multiple studies from different periods. Furthermore, *PHF19* has not been found to be significantly mutated in sequencing-based studies^28,29^, suggesting that its overexpression is not directly related to genomic alterations within the *PHF19* gene. We also show that a simple four feature predictor composed of age, ISS, and expression of *PHF19* and *MMSET* performs similarly to more complex models with many more gene expression features included (Supplemental Figure 4). This simplicity may allow the model to be more easily adopted in a clinical setting where only two genes would need to be measured.

Apart from its prognostic value, we show that *PHF19* has functional importance in MM. Knockdown of *PHF19* led to a significant reduction of growth and cell cycle arrest *ex-vivo*, suggesting that *PHF19* may play a role in transitioning cells into a highly proliferative state in MM. *PHF19* has been shown to be a major modulator of histone methylation thereby regulating transcriptional chromatin activity^30^. *PHF19* directly recruits the polycomb repressive complex 2 (PRC2) via binding to H3K36me3 and leads to activation of enhancer of zeste homolog 1 and 2 (EZH1/EZH2) as enzymatic subunits of PRC2, thereby resulting in tri-methylation of H3K27^31,32^. This process has been shown to enforce gene repression and is known to promote tumor growth in a variety of cancers^33^. While *MMSET* has also been shown to regulate histone methylation, its role as an epigenetic modulator is less well understood. Some reports have suggested that *MMSET* leads to transcriptional repression through generation of H4K20me^34^, H3K27me3^35^ or H3K36me3^35^, while other studies show that *MMSET* enhances transcription through generation of H4K20me2^36^ and H3K36me2^35^. In contrast to *MMSET, PHF19* expression is present in all MM subgroups and is preferentially overexpressed in high risk MM. These results are indicative of a strong correlation between increased histone methylation, in particular H3k27 trimethylation, and disease aggressiveness. Further work will be necessary to elucidate the mechanisms of *PHF19* in MM biology and any interplay with *MMSET*.

## Acknowledgements

Work completed by C.S. was funded by NCATS grant, KL2TR000063.

## Authorship and conflict-of-interest statements

C.S., V.L. and K.B. performed experiments; M.J.M., C.S., C.E., F.T., F.G., B.S.W., A.P. and E.C. analyzed results and made the figures; C.E., B.S.W., A.P., Y.G., H.C., Y.C., B.L., F.G., B.H. and M.O. built top performing and comparator models for predicting high-risk; T.Y., E.C.N., M.J.M., M.T., J.Gu. and B.S.W. designed and built compute infrastructure and model performance metrics for benchmarking methods; H.Y.D., W.S.D., H.G., H.A.-L., M.S., P.S., K.H.S., N.M., D.A., G.M., D.H. and B.A.W. provided key test datasets without which this investigation would not be possible; D.R., E.F., M.J.M., and K.M. processed and curated datasets; M.J.M., A.D., D.B., F.S., C.S., F.T., B.S.W., A.T., B.A.W., J.Gu. designed the research; M.J.M., A.D., C.S., F.T., B.S.W., A.R., S.A.D., A.T., B.A.W., J.Gu. wrote the manuscript. M.J.M, C.S., C.E., F.T, F.G. and A.D. contributed equally. D.B., J.G., B.A.W., A.T., J.Gu. contributed equally. A complete list of the members of the Multiple Myeloma DREAM Consortium and corresponding affiliations appears in the supplement text.

The following byline authors: F. T., K. M., K. B., F. S., E. F., A.R., S.A.D., A. D., D. B., A. T. have employment and/or equity ownership in Celgene. A.D. also had equity in Twinstrand Biosciences. H.G. has grants and/or provision of Investigational Medicinal Product: MM5 (Celgene, Janssen, Chugai), HD6 (BMS, Celgene, Chugai), HD7 (Sanofi, Celgene), MMRF (John Hopkins University), ReLApsE (Amgen, Celgene, Chugai, Dietmar-Hopp-Stiftung), research support (Amgen, BMS, Celgene, Chugai, Janssen, Molecular Partners, MSD, Sanofi, Mundipharma, Takeda and Novartis), advisory boards (Adaptive Biotechnology, Amgen, BMS, Celgene, Janssen, Sanofi and Takeda), honoraria (ArtTempi, BMS, Celgene, Chugai, Janssen, Novartis and Sanofi). N.M. has consulted for Abbvie, Adaptive, Amgen, Celgene, Janssen, Takeda and Oncopep and also has equity in Oncopep. K.S. is on advisory boards for: Celgene, BMS, Amgen, Takeda, Janssen and Sanofi Genzyme, has received research funding from AbbVie, and has consulted for Adaptive Biotechnologies.

## References

1. Palumbo A, Anderson K. Multiple myeloma. N Engl J Med. 2011;364(11):1046–1060.

2. Kyle RA, Therneau TM, Rajkumar SV, Larson DR, Plevak MF, Melton LJ 3rd. Incidence of multiple myeloma in Olmsted County, Minnesota: Trend over 6 decades. Cancer. 2004;101(11):2667–2674.

3. Kyle RA, Rajkumar SV. Multiple myeloma. N Engl J Med. 2004;351(18):1860–1873.

4. Sonneveld P, Avet-Loiseau H, Lonial S, et al. Treatment of multiple myeloma with high-risk cytogenetics: a consensus of the International Myeloma Working Group. Blood. 2016;127(24):2955–2962.

5. Jethava Y, Mitchell A, Zangari M, et al. Dose-dense and less dose-intense Total Therapy 5 for gene expression profiling-defined high-risk multiple myeloma. Blood Cancer J. 2016;6(7):e453.

6. Pawlyn C, Morgan GJ. Evolutionary biology of high-risk multiple myeloma. Nat Rev Cancer. 2017;17(9):543–556.

7. Shaughnessy JD Jr, Zhan F, Burington BE, et al. A validated gene expression model of high-risk multiple myeloma is defined by deregulated expression of genes mapping to chromosome 1. Blood. 2007;109(6):2276–2284.

8. Hervé A-L, Florence M, Philippe M, et al. Molecular heterogeneity of multiple myeloma: pathogenesis, prognosis, and therapeutic implications. J Clin Oncol. 2011;29(14):1893–1897.

9. Broyl A, Hose D, Lokhorst H, et al. Gene expression profiling for molecular classification of multiple myeloma in newly diagnosed patients. Blood. 2010;116(14):2543–2553.

10. Shi L, Campbell G, Jones WD, et al. The MicroArray Quality Control (MAQC)-II study of common practices for the development and validation of microarray-based predictive models. Nat Biotechnol. 2010;28(8):827–838.

11. Kryukov F, Nemec P, Radova L, et al. Centrosome associated genes pattern for risk sub-stratification in multiple myeloma. J Transl Med. 2016;14(1):150.

12. Dickens NJ, Walker BA, Leone PE, et al. Homozygous deletion mapping in myeloma samples identifies genes and an expression signature relevant to pathogenesis and outcome. Clin Cancer Res. 2010;16(6):1856–1864.

13. Meissner T, Seckinger A, Rème T, et al. Gene expression profiling in multiple myeloma--reporting of entities, risk, and targets in clinical routine. Clin Cancer Res. 2011;17(23):7240–7247.

14. Popovici V, Chen W, Gallas BG, et al. Effect of training-sample size and classification difficulty on the accuracy of genomic predictors. Breast Cancer Res. 2010;12(1):R5.

15. Attal M, Lauwers-Cances V, Hulin C, et al. Lenalidomide, Bortezomib, and Dexamethasone with Transplantation for Myeloma. N Engl J Med. 2017;376(14):1311–1320.

16. Laganà A, Perumal D, Melnekoff D, et al. Integrative network analysis identifies novel drivers of pathogenesis and progression in newly diagnosed multiple myeloma. Leukemia. 2018;32(1):120–130.

17. Went M, Sud A, Speedy H, et al. Genetic correlation between multiple myeloma and chronic lymphocytic leukaemia provides evidence for shared aetiology. Blood Cancer J. 2018;9(1):1.

18. Chattopadhyay S, Thomsen H, Yadav P, et al. Genome-wide interaction and pathway-based identification of key regulators in multiple myeloma. Commun Biol. 2019;2:89.

19. Boyd KD, Ross FM, Walker BA, et al. Mapping of chromosome 1p deletions in myeloma identifies FAM46C at 1p12 and CDKN2C at 1p32.3 as being genes in regions associated with adverse survival. Clin Cancer Res. 2011;17(24):7776–7784.

20. Wu P, Walker BA, Brewer D, et al. A gene expression-based predictor for myeloma patients at high risk of developing bone disease on bisphosphonate treatment. Clin Cancer Res. 2011;17(19):6347–6355.

21. Moreaux J, Rème T, Leonard W, et al. Development of gene expression-based score to predict sensitivity of multiple myeloma cells to DNA methylation inhibitors. Mol Cancer Ther. 2012;11(12):2685–2692.

22. Maes K, De Smedt E, Kassambara A, et al. In vivo treatment with epigenetic modulating agents induces transcriptional alterations associated with prognosis and immunomodulation in multiple myeloma. Oncotarget. 2015;6(5):3319–3334.

23. Kuiper R, Broyl A, de Knegt Y, et al. A gene expression signature for high-risk multiple myeloma. Leukemia. 2012;26(11):2406–2413.

24. Kassambara A, Gourzones-Dmitriev C, Sahota S, et al. A DNA repair pathway score predicts survival in human multiple myeloma: the potential for therapeutic strategy. Oncotarget. 2014;5(9):2487–2498.

25. Kuiper R, van Duin M, van Vliet MH, et al. Prediction of high- and low-risk multiple myeloma based on gene expression and the International Staging System. Blood. 2015;126(17):1996–2004.

26. Liptak T. On the combination of independent tests. Magyar Tud Akad Mat Kutato Int Kozl. 1958;3:171–197.

27. Ren Z, Ahn JH, Liu H, et al. PHF19 promotes multiple myeloma tumorigenicity through PRC2 activation. Blood. August 2019. doi:10.1182/blood.2019000578

28. Walker BA, Mavrommatis K, Wardell CP, et al. Identification of novel mutational drivers reveals oncogene dependencies in multiple myeloma. Blood. 2018;132(6):587–597.

29. Lohr JG, Stojanov P, Carter SL, et al. Widespread genetic heterogeneity in multiple myeloma: implications for targeted therapy. Cancer Cell. 2014;25(1):91–101.

30. Cai L, Rothbart SB, Lu R, et al. An H3K36 methylation-engaging Tudor motif of polycomb-like proteins mediates PRC2 complex targeting. Mol Cell. 2013;49(3):571–582.

31. Brien GL, Gambero G, O’Connell DJ, et al. Polycomb PHF19 binds H3K36me3 and recruits PRC2 and demethylase NO66 to embryonic stem cell genes during differentiation. Nat Struct Mol Biol. 2012;19(12):1273–1281.

32. Ballaré C, Lange M, Lapinaite A, et al. Phf19 links methylated Lys36 of histone H3 to regulation of Polycomb activity. Nat Struct Mol Biol. 2012;19(12):1257–1265.

33. Yamagishi M, Uchimaru K. Targeting EZH2 in cancer therapy. Curr Opin Oncol. 2017;29(5):375–381.

34. Marango J, Shimoyama M, Nishio H, et al. The MMSET protein is a histone methyltransferase with characteristics of a transcriptional corepressor. Blood. 2008;111(6):3145–3154.

35. Nimura K, Ura K, Shiratori H, et al. A histone H3 lysine 36 trimethyltransferase links Nkx2-5 to Wolf-Hirschhorn syndrome. Nature. 2009;460(7252):287–291.

36. Kang H-B, Choi Y, Lee JM, et al. The histone methyltransferase, NSD2, enhances androgen receptor-mediated transcription. FEBS Lett. 2009;583(12):1880–1886.

