## Supplemental Text for "Multiple Myeloma DREAM Challenge Reveals Epigenetic Regulator *PHF19* As Marker of Aggressive Disease"

**Supplemental Methods**

**Cohort Identification**

*Training datasets*: The Multiple Myeloma DREAM organizers identified over 40 publicly available myeloma datasets via NCBI’s Gene Expression Omnibus (GEO) and European Bioinformatics Institute Array Express (EMBL-EBI) databases. These datasets were then filtered to ensure that they contained newly diagnosed patients, progression and survival outcome, CD138 positive samples, and unique samples not contained in other studies. Three microarray datasets passed filtering and were included as training sets in this challenge: HOVON-65/GMMG-HD4 ^1^, GSE24080 (University of Arkansas Medical Sciences) ^2,3^, E-MTAB-4032 (Masaryk University) ^4^. The Multiple Myeloma Research Foundation (MMRF) provided an additional training dataset from its CoMMpass study (release IA9) ^5^, which included RNA-seq expression, variant and clinical outcome data from newly diagnosed myeloma patients.

Validation cohorts: Potential validation datasets must have clinical outcome data that has never been made publically available. The search of GEO and EMBL-EBI identified multiple datasets that passed the above filters but with clinical data never released. To obtain the missing clinical outcome data, we independently contacted the corresponding authors of two studies, E-MTAB-372 (Heidelberg University)^6^ and MRC Myeloma IX (MRC-IX) ^7^. They agreed to make clinical outcome data available for the purposes of this Challenge thus enabling these datasets to be used as validation cohorts. Collaborators in the Myeloma Genome Project provided two private datasets from clinical trials: the MGP-UAMS dataset from the Myeloma IX trial ^8^ comparing standard chemotherapy regimen (cyclophosphamide, dexamethasone plus thalidomide) to an experimental arm (cyclophosphamide, dexamethasone plus lenalidomide with or without carfilzomib), and the MGP-DFCI dataset wherein standard dose lenalidomide plus bortezomib and dexamethasone (RVD) followed by lenalidomide maintenance was compared to an intensive treatment pathway of RVD followed by high dose therapy and autologous stem cell transplant then followed by lenalidomide maintenance ^9,10^. Importantly, the intensive and non-intensive treatment pathways were not assigned based on patient age; instead they were randomized. An additional dataset from the Oncology Research Information Exchange Network (ORIEN) was made available through a collaboration with Moffitt Cancer Center and M2Gen.

*Patient Populations:* All datasets are comprised of samples of CD138 positive plasma cells isolated from bone marrow aspirate or blood of newly diagnosed patients. Additionally, patient gender, age and International Staging System (ISS) classification are provided in each dataset, though may be missing for individual samples. Many datasets also include cytogenetic calls for common translocations and deletions in MM. For dataset summary statistics see table 1.

**Data Curation**

Clinical data across all cohorts was standardized according to a common clinical dictionary provided by MMRF. Progression events based on FDA definitions were adjusted to resemble the European Medicines Agency definition of progression, i.e., with any death classified as a progression event. For datasets lacking ISS , we reached out to data originators for sample beta 2-microglobulin and albumin serum levels and computed ISS accordingly. Expression microarrays were processed with the R “oligo” package. RNA-seq and WES data were processed to match the TGEN pipeline. Specifically, gene expression was calculated by processing raw RNA-seq FASTQ files with Salmon^11^, compressing transcripts to hg19 genes via the tximport package^12^.

**Estimated cytogenetics**

Molecular data refers to assays measuring cytogenetics: chromosomal deletions, translocations, and duplications. Many studies included here have FISH (fluorescent in situ hybridization) targeting specific regions which are often cytogenetically modified in MM. The MMRF training dataset has Seq-FISH which is WGS used to quantify deletions, translocations, and duplications.

The MMRF data and validation data with sequencing data (UAMS Whole Exome, Dana-Farber Cancer Institute RNA-seq) also have expert-curated calls for common cytogenetic features that take into account FISH quality, ploidy and other factors. The expert curated cytogenetic labels are blinded for the purposes of the challenge [i.e., the features are simply identified as feature 1, feature 2, ..., feature 18, rather than as t(4;14), del(17p), etc.]. Those expert-curated calls can be used in modeling for challenges 1 and 3. We recommended against using raw FISH or seqFISH based calls from MMRF in modeling given that only the expert-curated calls will be available for the corresponding validation cohorts.

**Challenge Procedures and Infrastructure**

The Challenge was hosted on Sage Bionetwork’s cloud-based data-sharing platform, Synapse, which provided access to training data, model submission, myeloma background information and other resources. In contrast to a traditional "data to model" framework in which challenge participants have direct access to blinded validation data, we used a "model to data" framework, where participants bundle their predictor with necessary software libraries and operating system into a “container,” which is then run in a cloud environment where the validation data is hosted^13^ (Figure 1). In this Challenge containerizaton was implemented via Docker^14^ (see supplement for additional details). Using this “model to data” approach allowed us to sequester validation data in the cloud and satisfied data provider requirements that data could not be shared directly with participants as the data was proprietary or pre-publication. As such, we were able to double the size of the validation cohorts for all three sub-Challenges. As an additional benefit, the “containerized” models allows for simplified dissemination to the research community, and for evaluation on future myeloma datasets.

The challenge was executed over three phases: an open phase, a leaderboard phase and validation phase. In the open phase, participants were unable to submit models for scoring but had access to training data, the challenge forum, challenge express queues in which models are run on test data to check for proper output formatting but are not scored. This open phase enabled participants to get accustomed to data formats and learn Docker, train models and post relevant questions to the community. In the leaderboard phase teams submitted their Dockerized algorithms, output from them was checked for proper prediction formatting and their accuracy was assessed on a subset of the validation data called the leaderboard data using the scoring metrics. This dataset was set aside to allow teams to assess and improve their models. Finally, in the validation phase predictors are assessed on the remainder of the validation cohort and ranked by accuracy.

The leaderboard phase was run using 20% of their respective validation data. The leaderboard phase had three rounds, in which participants were given access to data bootstrapped from the original 20%. They were provided feedback on the performance of up to 2 submitted models. The remaining 80% of the validation set was used in the validation round where participants were allowed one model submission to determine their final score.

**Model Scoring**

Participant prediction models had to provide both continuous estimates of risk for each patient and a binary high-risk call defined as progressing prior to 18 months. To identify top performing teams we employed two metrics to assess the accuracy of submitted models within a given validation cohort: the integrated AUC (iAUC) and balanced accuracy (BAC). While the area under the ROC curve (AUC) is a widely accepted metric of prediction accuracy, it is sensitive to the specific threshold used to define patient risk. The myeloma research community has not yet reached consensus on the time point that best separates patients into risk groups though there is general agreement that it lays somewhere between 1 and 2 years. Thus we chose the more robust iAUC ranging from 12 to 24 months sampled at weekly intervals as the primary metric. iAUCs computed in each cohort were combined into a weighted average (wAUC) with each cohort iAUC weighted by the square of the number of high-risk patients in it.

We used the Bayes factor, *K*, to identify statistically tied top performing predictors in a given sub-challenge. We used 1000 bootstrapped samples of the validation dataset to compute the distribution of each model’s iAUC. *N_i_* observations were sampled with replacement from each validation cohort, where *N_i_* is the number of observations in cohort *i*. These distributions are then used to compute the Bayes factor *K*_p_ for each high-risk prediction model *p* with:

$K_{p}=\frac{\sum_{j=1}^{1000} {wiAUC}_{p,j} < {wiAUC}_{best,j}}{\sum_{j=1}^{1000} {wiAUC}_{p,j} \geq{wiAUC}_{best,j}}$

Where *best* simply indicates the model with the highest wAUC prior to bootstrapping. Predictors with *K_p_< 3* are considered tied with the *best* model and the weighted BAC was used as a tie-breaking metric in order to determine the top performing model among them. The BAC was used since it is based on a thresholded call of high-risk as would likely be the case of any model adapted for clinical use. Similar to the wAUC, the weighted BAC’s were computed in each validation cohort and then averaged using the square of the number of high-risk patients in each dataset.

**Comparator Models**

Throughout the leaderboard and validation phases, challenge organizers submitted four types of comparator models to which challenge participant’s predictors were compared: baseline models, previously-published myeloma risk models, combination models and an ensemble model which had access to an indicator of whether a patient received an autologous stem cell transplant. The baseline predictors were Cox proportional hazard models where progression free survival was modeled on age or ISS or age and ISS together. As comparators these models represent a reasonable lower bound on expected performance of high-risk classifiers.

Additionally, challenge organizers implemented versions of the popular microarray based UAMS GEP-70 ^15^ and EMC-92^16^ predictors, where model coefficients were adjusted to use gene-level expression instead of probe-level expression. Finally, we extended these models to include age, ISS or both.

Comparator models are implemented in the following directories in the GitHub repository at (<https://github.com/Sage-Bionetworks/Celgene-Multiple-Myeloma-Challenge/tree/master/docker-images>):

| Model | Directory |
| --- | --- |
| Age | age-coxph |
| ISS | iss-coxph |
| Age + ISS | age-iss-coxph |
| UAMS70 | uams-70-sc2-and-3 |
| UAMS70 + Age | uams-70-age-sc2-and-3 |
| UAMS70 + ISS | uams-70-iss-sc2-and-3 |
| UAMS70 + Age + ISS | uams-70-age-iss-sc2-and-3 |
| EMC92 + Age | emc-92-age-sc2-and-3 |
| EMC92 + ISS | emc-92-iss-sc2-and-3 |
| EMC92 + Age + ISS | emc-92-age-iss-sc2-and-3 |

The HOVON-65/GMMG-HD4 was not used for training any comparator model that included age as a covariate, as age was not included in this datasets’s clinical annotations. When included as a covariate, ISS was represented as a categorical variable (factor) not an integer. Missing clinical annotations for ISS, Gender, and Age were imputed by using the mode (i.e., most commonly-occurring value) within each study.

Baseline Models: age, ISS, and age + ISS

Baseline models were Cox proportional hazards models where PFS was the response variable and age, ISS, or both age and ISS were explanatory variables. Continuous risk predictions scores of each sample were thresholded to create high risk classifications. The threshold of a given model was calculated by generating sample level predictions in the training data. The threshold was defined as the cutpoint that maximized the logrank statistic comparing above and below the threshold to their true high-risk status. Continuous predictions for validation data were computed by applying the above trained Cox proportional hazards model to them. These values were dichotomized into high and low risk according to whether the prediction was above or below, respectively, the threshold.

Published UAMS70 and EMC92 models:

The UAMS70 and EMC92 models are based on published implementations [PMID: 22722715]. In both cases, the models were trained by: (1) translating the published Affymetrix HG U133 Plus 2.0 probesets for the respective model to Entrez identifiers; (2) defining gene-level coefficients by adjusting published probeset-level coefficients to account for cases in which a single probeset maps to multiple genes; (3) averaging biological replicates (i.e., multiple samples from the same patient) to define a single per-patient expression data; (4) restricting genes applied in the model to the intersection of the published gene set and genes provided in the validation data; (5) standardizing genes independently within each dataset (i.e., such that each gene has mean zero and standard deviation one within each dataset); (6) computing a continuous score as the sum of the (log2-normalized) gene expression values, weighted by the gene-level coefficients; and (7) defining an optimal threshold as a cutpoint of these continuous scores using the logrank statistic, as described in the baseline models section above.

Probeset identifiers were translated to Entrez identifiers using the getBM function from R library biomaRt, passing it arguments attributes=c(‘affy_hg_u133_plus_2’, ‘entrezgene’), filters=’affy_hg_u133_plus_2’, and mart=useMart(“ensembl”,dataset=”hsapiens_gene_ensembl”).

The UAMS70 model defines a score as the mean expression of the “down” probes subtracted from the mean expression of the “up” probes. Hence, the probeset-level weights (i.e., prior to adjustment) are simply 1/*n_up_* and 1/*n_down_* for the “up” and “down” probes, respectively, where *n_up_* is the number of “up” genes and *n_down_* is the number of “down” genes. Probeset-level EMC92 coefficients were taken from the published implementation.[PMID: 22722715]

The probeset-level weight (or coefficient) *W_p_* associated with a probeset *P* that is mapped to the *n* genes *g_i_* (*i* = 1 … *n*) was adjusted to the gene-level weight *W_g_* = *W_p_* / *n*.

Continuous predictions for validation data were computed using steps (1) - (6) described above. These values were dichotomized into high and low risk according to whether the prediction was above or below, respectively, the threshold defined in step (7) above.

Extended Models:

Each extended model is a Cox proportional hazards model where the response variable was PFS and the explanatory variables include the raw, continuous score output from the published UAMS70 or EMC92 model described above and age, ISS, or age and ISS as described in the baseline models. For example, the UAMS70 + age + ISS model is a Cox proportional hazards model with age, ISS, and the raw, continuous UAMS70 score as explanatory variables. The continuous score of the extended model is the prediction of the Cox proportional hazards model. As described above, a threshold is defined using the logrank statistic during training and applied during validation. Data were pre-processed as described above in steps (1) - (6) for both training and validation.

**The Internal Collaborator Comparator Model (REFS^TM^) with ASCT Indicator**

Working with GNS healthcare we used their Reverse-Engineering and Forward Simulation method (REFS, (Latourelle et al. 2017) to create an internal comparator model (henceforth referred to as the REFS model). The REFS predictor focused on logistic modeling of the 18-month binarized PFS outcome.

In addition to other clinical features, the REFS model used an indicator for a patient undergoing autologous stem cell transplant status (ASCT). While ASCT status was not included in the clinical files explicitly provided to challenge participants, all ASCT status annotation for training data was publicly available elsewhere, and we had further access to it for part of the validation data (that from DFCI). Since this was the only model with theoretical access to the DFCI annotation during model development, we treated the REFS model as an upper-bound comparator, although this data was not actually used by our team in any way.

The REFS engine permits construction of large generalized linear model ensembles sampled from overall model space consistent with training data. It enables model ensembles of 1000s of models sampled from 50000-variable space, while incorporating multiple data modalities and modeling up to 3-way interactions. The sampling nature of the optimizer, as well as the ensemble approach to modeling, permit accurate inference over very under-determined datasets.

The REFS engine’s ensemble approach enables the importance of a given clinical feature or gene to be assessed through cross-validation and feature perturbation. The importance of each variable was estimated using a procedure similar to the one that defines the *predictive effect size* ([Hu and Ishwaran, 2017](https://arxiv.org/abs/1701.04944)). The training dataset was split using repeated k-fold cross-validation (specifically, 5 folds repeated 5 times) and used to calculate the average iAUC over folds and repeats. This is the reference out-of-sample performance of the model. Then the observed values of one of the variables were permuted and the predictions and average iAUC were recalculated. The difference between the reference performance and the new performance after permuting the variable defined the importance of that variable. This procedure is then repeated for all variables in the model.

**Top Performing Method (Genome Institute of Singapore)**

*Summary Sentence*

We combined sets of genes associated with Multiple Myeloma pathways and chromosomal abnormalities in a systems approach and iteratively selected the most informative features for classification of high-risk and low-risk Multiple Myeloma patients using regularized regression [1].

*Background/Introduction*
Multiple Myeloma (MM) is a disease that arises from a complex set of genetic changes. The genes and chromosomal regions that are affected can differ substantially between patients, making Multiple Myeloma a genetically highly heterogeneous disease. It was observed that a subset of MM patients had poor survival outcome despite aggressive treatment, which may be partially explained through the genetic and molecular characteristics of the disease. However, due to its heterogeneity and the large number of possible features (genes), using expression of individual genes may not be sufficient to accurately and robustly distinguish between high- and low-risk patients. Interestingly, even diverse genetic changes can have similar effects downstream, for example by altering the activity of the same pathway through distinct mechanisms, or the over- or under-expression of genes resulting from amplification or deletion of a chromosomal region. This suggests, that the heterogeneity of the genetic alterations can partially be controlled by using supersets of genes that are expected to result in similar molecular phenotypes. There has been research on the mutations and pathways associated with Multiple Myeloma over the years, with several risk markers developed based on gene expression profiles [2, 3]. Here, we have adopted a systems approach to combine sets of genes in pathways and chromosomal abnormalities that have been implicated in high-risk Multiple Myeloma or that have similar molecular functions. This approach handles the underlying genetic heterogeneity and at the same time enables a reduction of the feature space through compression, thereby enabling us to train a robust, generalisable model. Together we engineered 28 features and trained a model using regularized logistic regression that achieved reproducibly good accuracy across different datasets on cross-validation for binary classification of high- and low-risk Multiple Myeloma.

*Methods*
For the gene expression-based predictor of high-risk Multiple Myeloma in sub-challenge 2, both the microarray and RNA-Seq expression data were combined into a training dataset. The RNA-Seq data was first normalized using log transformation, and both the microarray and RNA-Seq data were next standardized using z-scores for all genes.
We next performed feature engineering to create sets of genes mapped to certain pathways and chromosomal abnormalities implicated in Multiple Myeloma, as well as other gene expression profiles developed by others, which includes:

I. Chromosomal abnormalities [3]: deletion of chromosome 1p, gain of chromosome 1q, gain of chromosome 9, deletion of chromosome 13q, deletion of chromosome 17p, translocation t(4;14), trainslocation t(11;14), translocation t(14;16/14;20).
II. DNA repair pathways [5]: non-homologous end-joining pathway, homologous recombination pathway, Fanconi anemia pathway, nucleotide excision repair pathway, mismatch repair pathway, base excision repair pathway.
III. Other pathways [3]: cell cycle pathway, p53 signalling pathway, NF-kB signalling pathway, Ras-ERK pathway.
IV. Genes targeted by Multiple Myeloma treatments [6]: Bortezomib, Thalidomide
V. Mutations associated with high-risk Multiple Myeloma [3].
VI. Other gene expression profiles obtained from literature: EMC92 [7], UAMS70 [8], DNA repair pathway score [4], IFM group [9], cell death network [10].

Sum of gene expressions were used, and feature selection of the engineered features was also performed to progressively discard uninformative features. We also include clinical data comprising age and ISS as features.


Samples with missing data were discarded for the training dataset, and only genes that were found across all training and validation datasets were retained. To minimize the problem of class imbalance in training, random undersampling was performed to ensure equal class distribution between high-risk and low-risk samples in the training dataset. The machine learning algorithm chosen for the construction of the final prediction model was regularized logistic regression, implemented using the glmnet package[11] in R. Five-fold cross-validation was simulated 100 times to estimate prediction model performance. The final model was comprised of one regularized logistic regression model trained on a balanced training dataset, classifying between high-risk and low-risk Multiple Myeloma.

*Conclusion/Discussion*
Multiple Myeloma is a complex disease, resulting from a multi-step progression of different mutations and chromosomal abnormalities. It is therefore difficult to use single mutations or gene expression to characterize high-risk Multiple Myeloma due to the different molecular and cytogenetic profiles even among high-risk patients. Adopting a systems approach by taking into account the gene expressions in pathways and chromosomal abnormalities associated with Multiple Myeloma enabled us to build a more robust model compared to single gene features leading to increased predictive performance of high-risk Multiple Myeloma.

**Second Place Feature Selection Methods**

The second place team, SUGO, used a simple univariate approach to rank gene for feature selection in building expression based models. In each of the four training datasets they computed each gene’s effect size, *z*, via the concordance index between overall survival and the gene’s expression. These effect sizes were then combined across training sets using Stouffer’s method with no weighting to yield a single meta-z per gene. They employed this meta concordance index method under two expression normalization procedures with CDKN3 and PHF19 coming out on top respectively and combined them with clinical features to create their model. For more information on SUGO’s model please see their full description in Synapse (<https://www.synapse.org/#!Synapse:syn10380508/wiki/499377>)

**Software**

R version 3.3.1 was used for data curation and statistical analyses in this DREAM Challenge. Specific packages used for curation and analysis: caret version 6.0-79, data.table 1.10.4-3, oligo 1.38.0, org.Hs.eg.db 3.4.0, plyr 1.8.4, pROC 1.11.0, risksetROC 1.0.4, survival 2.38-3, synapseClient 1.13-4, timeROC 0.3. The top-performing model also used biomaRt, caTools, dplyr, FSelectorm, glmnet 2.0-5, missForest and scales. Docker version 17.05.0 was used for challenge submission and running containers. Challenge documentation, including a detailed description of its design, overall results, scoring scripts, data dictionary, and training data along with participant methods and code can be accessed via the Synapse platform at [synapse.org/MultipleMyelomaChallenge](http://www.synapse.org/MultipleMyelomaChallenge).

**Functional Assays**

*Cell lines and culture*

The multiple myeloma cell lines JJN3 and ARP1 were grown in RPMI medium supplemented with 10% fetal bovine serum and 1% penicillin/streptomycin.

*Lentiviral short hairpin RNA vectors and transduction*

*PHF19* targeting short hairpin RNA (shRNA) with tetracycline controlled transcriptional activation (TRIPZ) was used for knockdown studies (Clone ID: V2THS_21282, Dharmacon). TRIPZ inducible lentiviral non-silencing shRNA was used as a negative control (Dharmacon). For production of lentiviral particles, lentiviral shRNA expression constructs were transfected together with packaging vectors into 293T producer cells using Lipofectamine LTX transfection reagent (Invitrogen), and supernatants were harvested after 48 and 72 hours and concentrated by ultracentrifugation. The JJN3 and ARP1 MM cell lines were transduced with the shRNA-containing lentiviruses and cultured in fresh medium. Puromycin at a concentration of 1µg/ml was added 72 hours after transduction to select for transduced cells. After 2 weeks of puromycin selection, doxycycline (1µg/ml) was added to the cells to induce the production of *PHF19* knockdown (KD) and control shRNA. *PHF19* KD efficiency was measured by real-time PCR and Westernblotting after 72 hours (JJN3) and 168 hours (ARP1) of doxycycline induction as described below.

*Real time polymerase chain reaction*

RNA was extracted with an RNeasy Micro kit (Qiagen), and quantity was assessed with the Qubit 4 Fluorometer (Thermo Scientific). Extracted RNA was reverse-transcribed with Superscript III reverse transcriptase (Invitrogen) and expression of *PHF19* was measured via Taqman assay (Hs01106991_m1, Thermo Scientific) as per manufacturer’s description (Applied Biosystems).

*Cell lysis and Westernblotting*

Cells were lysed in lysis buffer (Thermo Scientific) and Westernblotting was performed as previously described using a *PHF19* directed antibody (Milipore Sigma) ^1^.

*Cell viability assay*

JJN3 and ARP1 cell lines containing the *PHF19* KD or control plasmid were incubated with 1µg/ml doxycycline for 72hrs (JJN3) or 168 hrs (ARP1) to allow for induction of *PHF19* KD and control shRNAs and then plated at 5x10^3^ cells per 100/µL. Viability was assessed by addition of Cell Titer Blue (Promega) and measured via Fluostar Omega Microplate reader (BMG Labtech) at various time points.

*Cell cycle analysis*

After 72hrs (JJN3 cell line) or 168hrs (ARP1) of doxycycline induction (1µg/ml), 1x10^6^ *PHF19* KD and control cells were washed and incubated with Vybrant DyeCycle Stain (Thermo Scientific) as per manufacturer’s description. Cell cycle was analyzed by flow cytometry using a FACSVerse flow cytometer (BD Biosciences) as previously described ^2^.

*Apoptosis*

To determine viability after *PHF19* KD, 1x10^6^ PHF19KD and control cells were washed and incubated with FITC conjugated Annexin V (BioLegend) and 7AAD after 72hrs (JJN3) or 168hrs (ARP1) of doxycycline induction (1µg/ml). The percentage of alive, apoptotic and necrotic cells was analyzed by flow cytometry using a FACSVerse flow cytometer (BD Biosciences) as previously described ^2^.

References

1.	Broyl A, Hose D, Lokhorst H, et al. Gene expression profiling for molecular classification of multiple myeloma in newly diagnosed patients. *Blood*. 2010;116(14):2543–2553.

2.	Shi L, Campbell G, Jones WD, et al. The MicroArray Quality Control (MAQC)-II study of common practices for the development and validation of microarray-based predictive models. *Nat. Biotechnol.* 2010;28(8):827–838.

3.	Popovici V, Chen W, Gallas BG, et al. Effect of training-sample size and classification difficulty on the accuracy of genomic predictors. *Breast Cancer Res.* 2010;12(1):R5.

4.	Kryukov F, Nemec P, Radova L, et al. Centrosome associated genes pattern for risk sub-stratification in multiple myeloma. *J. Transl. Med.* 2016;14(1):150.

5.	Miller A, Cattaneo L, Asmann YW, et al. Correlation Between Somatic Mutation Burden, Neoantigen Load and Progression Free Survival in Multiple Myeloma: Analysis of MMRF CoMMpass Study. *Blood*. 2016;128(22):193–193.

6.	Meissner T, Seckinger A, Rème T, et al. Gene expression profiling in multiple myeloma--reporting of entities, risk, and targets in clinical routine. *Clin. Cancer Res.* 2011;17(23):7240–7247.

7.	Dickens NJ, Walker BA, Leone PE, et al. Homozygous deletion mapping in myeloma samples identifies genes and an expression signature relevant to pathogenesis and outcome. *Clin. Cancer Res.* 2010;16(6):1856–1864.

8.	Morgan GJ, Davies FE, Gregory WM, et al. First-line treatment with zoledronic acid as compared with clodronic acid in multiple myeloma (MRC Myeloma IX): a randomised controlled trial. *Lancet*. 2010;376(9757):1989–1999.

9.	Walker BA, Samur MK, Mavrommatis K, et al. The Multiple Myeloma Genome Project: Development of a Molecular Segmentation Strategy for the Clinical Classification of Multiple Myeloma. *Blood*. 2016;128(22):196–196.

10.	Attal M, Lauwers-Cances V, Hulin C, et al. Lenalidomide, Bortezomib, and Dexamethasone with Transplantation for Myeloma. *N. Engl. J. Med.* 2017;376(14):1311–1320.

11.	Patro R, Duggal G, Love MI, Irizarry RA, Kingsford C. Salmon provides fast and bias-aware quantification of transcript expression. *Nat. Methods*. 2017;14(4):417–419.

12.	Soneson C, Love MI, Robinson MD. Differential analyses for RNA-seq: transcript-level estimates improve gene-level inferences. *F1000Res.* 2015;4:1521.

13.	Guinney J, Saez-Rodriguez J. Alternative models for sharing confidential biomedical data. *Nat. Biotechnol.* 2018;36(5):391–392.

14.	Boettiger C. An introduction to Docker for reproducible research. *Oper. Syst. Rev.* 2015;

15.	Shaughnessy JD Jr, Zhan F, Burington BE, et al. A validated gene expression model of high-risk multiple myeloma is defined by deregulated expression of genes mapping to chromosome 1. *Blood*. 2007;109(6):2276–2284.

16.	Kuiper R, Broyl A, de Knegt Y, et al. A gene expression signature for high-risk multiple myeloma. *Leukemia*. 2012;26(11):2406–2413.

**Supplemental Figure Legends**

Supplemental Figure 1, Feature selection in GIS’s top performing model: Mean decrease in Gini coefficient from a random forest classifier was used to determine feature importance of public classifiers and clinical measures. Blue indicates features that were then included in penalized classifier.

Supplemental Figure 2, Feature importance when adding *PHF19* as a possible covariate in GIS’s top performing: Mean decrease in Gini coefficient from a random forest classifier. Blue indicates features that were then included in penalized classifier. *PHF19* is noted in green and was also selected for the penalized classifier.

Supplemental Figure 3, Assessment of the effect of *PHF19* Knock down on cell cycle and apoptosis in JNN3 and ARP1 cell lines: A) Representative density plot of the JJN3 cell line shows increased amount of cells in the G0/G1 stage after PHF19 knockdown compared to scrambled control. B) Repeated PHF19 knock down experiments in JJN3 cell line (5 control and 5 KD) and ARP1 cell line (4 control and 4 KD) demonstrate significantly increased G0/G1 cell cycle arrest after. Significance assessed by the t-test of a PHF KD indicator coefficient in a linear regression model (*M-G2% ~ cell line + PHF19 KD* p-value calculated by PHF19KD coefficient). The model included cell line to control for its effect on cell cycle. Analysis of apoptosis/necrosis with annexin V/DAPI in JJN3 and ARP1 cells transfected with scrambled control or *PHF19 KD* shRNAs did not show any significant difference between the control and *PHF19 KD* groups. Panel C shows FACS contour plots of 1 representative experiment. Panel D and E show the mean +/- SD of 3 independent experiments in JJN3 (D) and ARP1 (E) cells.

Supplemental Figure 4, Predictions from simple four feature classifier are similar to UAMS-70 extended predictions: continuous prediction scores from the UAMS-70 model extended to include age and ISS show a strong linear relationship with prediction scores from the age+ISS+PHF19+MMSET model in all four validation cohorts.

Authors Consortium

T Aderinwale^29^, T Afonso^30^, A Agibetov^31^, P Agrawal^32^, C Andresen^33,34^, S Bae^35,36^, J Bao^37^, S Bhalla^38^, K Boroevich^39^, B Brors^40,41^, J Causey^42^, K Chaudhary^43^, J Cheng^44^, Y Choi^45^, S Correia^30^, J Cursons^46^, H Demirci^30^, A DEYATI^47^, S Dhanda^48^, W Dong^49^, L Elo^50^, W Fang^51^, Q Feng^52^, P Ferreira^53^, M Foroutan^54^, J Gagnon-Bartsch^55^, P Georgeson^56^, J Göke^37^, C Guziolowski^57^, Z Han^58^, H Hassanzadeh^59^, N Hawkins^60^, e heo^36,61^, C Hong^40^, X Huang^42^, K Huang^58^, D Huebschmann^33,34^, M Hwang^51^, C Imbusch^40^, M Jeon^45^, J Ji^62^, S Jiang^63^, T Johnson^58,64^, S Jung^65^, K Kang^66^, J Kang^45,67^, H Kaur^32^, H Kazan^68^, D Kesar^69^, D Kim^45^, S Kim^45^, K Kim^45^, Y Kim^70^, S Kim^70^, R Klén^50^, A Kodumuri^71^, R Kurilov^40^, C Kurz^72,73^, M Larmuseau^74,75^, b lee^76^, H Lee^67^, M Lee^70^, T Lee^70^, A Lima^30^, R Luethy^77^, A Lysenko^39^, T Ma^78^, R Madduri^65,79^, S Madduru^80^, S Mahajan^48^, M Mahmoudian^81^, K Marchal^74,75^, J Markham^82^, A McGlinchey^83^, B Miannay^57^, R Molania^84^, M Mushthofa^74,75^, G Narasimhan^85,86^, B Panwar^48^, s park^76^, X Pastor^33^, G Paternostro^87^, L Peixoto^30^, S Piccolo^88^, C Piermarocchi^60^, J Qualls^42^, G Raghava^89^, M Rocha^30^, R Rodrigues^30^, H Ryu^90^, M Samwald^31^, S Santos^30^, A Sathe^91^, F Seyednasrollah^50^, H Shih^51^, L Sieverling^40^, N Smith^92^, T Speed^46^, T Szedlak^60^, T Tsunoda^39,93^, S Usmani^32^, D Van Daele^94^, V Vieira^30^, J Wang^95^, Q Wang^40^, b weytjens^96,97^, S Wong^40^, C Wu^51^, Y Wu^51^, J Xu^40,98^, C Yu^58,64^, J Zhang^99^

29. Electrical and Computer Engineering Graduate Program, Antalya Bilim University, Antalya, Turkey
30. Centre of Biological Engineering, Universidade do Minho, Braga, Braga, Portugal
31. Section for Artificial Intelligence and Decision Support, Medical University of Vienna, Vienna, Austria
32. Bioinformatics Center, CSIR-Institute of Microbial Technology, Chandigarh, Chandigarh, India
33. Theoretical Bioinformatics, German Cancer Research Center (DKFZ), Heidelberg, Germany
34. Pattern Recognition and Digital Medicine, Heidelberg Institute for Stem Cell Technology and Experimental Medicine, Heidelberg, Germany
35. Department of Biological Science, Department of Bio-Brain Engineering, Korea Advanced Institute of Science and Technology, Daejeon, South Korea
36. Deargen Inc., Daejeon, South Korea
37. Computational and Systems Biology, Genome Institute of Singapore, Singapore, Singapore
38. Department of Computational Biology, Indraprastha Institute of Information Technology, Delhi, India
39. Center for Integrative Medical Sciences, RIKEN, Yokohama, Kanagawa, Japan
40. Applied Bioinformatics, German Cancer Research Center (DKFZ), Heidelberg, Germany
41. German Cancer Consortium (DKTK), Heidelberg, Germany
42. Department of Computer Science, Arkansas State University, Jonesboro, AR, USA
43. Epidemiology Program, University of Hawaii Cancer Center, Honolulu, Hawaii, USA
44. School of Biomedical Engineering, Shenzhen University, Shenzhen, China
45. Department of Computer Science and Engineering, Korea University, Seoul, South Korea
46. Bioinformatics Division, Walter & Eliza Hall Institute of Medical Research, Parkville, Victoria, Australia
47. Bioinformatics, Biocon Bristol-Myers Squibb R&D Center, Bangalore, Karnataka, India
48. Division of Vaccine Discovery, La Jolla Institute for Immunology, La Jolla, California, USA
49. Ann Arbor Algorithms Inc., Ann Arbor, Michigan, USA
50. Turku Bioscience Centre, University of Turku and Åbo Akademi University, Turku, Finland
51. Institute of Biomedical Sciences, Academia Sinica, Taipei, ROC
52. School of Biomedical Engineering, Southern Medical University, Guangzhou, China
53. IPATIMUP/i3S and Faculty of Sciences, University of Porto, Porto, Porto, Portugal
54. Department of Clinical Pathology, The University of Melbourne Centre for Cancer Research, Victorian Comprehensive Cancer Centre, The University of Melbourne, Melbourne, Victoria, Australia
55. Department of Statistics, University of Michigan, Ann Arbor, Michigan, USA
56. Faculty of Medicine, Dentistry and Health Sciences, The University of Melbourne, Melbourne, Victoria, Australia
57. Laboratoire de Sciences du Numérique de Nantes (LS2N), Ecole Centrale de Nantes
58. Department of Medicine, Indiana University School of Medicine, Indianapolis, IN, USA
59. School of Computational Science and Engineering, Georgia Institute of Technology, Atlanta, Georgia, USA
60. Department of Physics and Astronomy, Michigan State University, East Lansing, Michigan, USA
61. School of Computing, Korea Advanced Institute of Science and Technology, Daejeon, South Korea
62. School of Statistics, Shandong University of Finance and Economics, Jinan, Shandong, China
63. Department of Computer Science, The University of Chicago, Chicago, IL, USA
64. Department of Biomedical Science, The Ohio State University, Columbus, OH, USA
65. Globus, University of Chicago, Chicago, IL, USA
66. Division of platform development, Deargen Inc., Daejeon, South Korea
67. Interdisciplinary Graduate Program in Bioinformatics, Korea University, Seoul, South Korea
68. Department of Computer Engineering, Antalya Bilim University, Antalya, Turkey
69. Department of computer science and engineering, Indraprastha Institute of Information Technology, Delhi, Delhi, India
70. Department of Computer Science and Engineering, Chung-Ang University, Seoul, South Korea
71. Mathematics and Computer Science, Argonne National Laboratory, Lemont, IL, USA
72. Institute of Health Economics and Health Care Management, Helmholtz Zentrum Muenchen, Neuherberg, Germany
73. Division of Pharmacoepidemiology and Pharmacoeconomics, Department of Medicine, Brigham and Women’s Hospital and Harvard Medical School, Boston, MA, USA
74. Department of Plant Biotechnology and Bioinformatics, Ghent University, Ghent, Ghent, Belgium
75. Department of Information technology (IDLab), Ghent University, Ghent, Belgium
76. Division of A.I. research, Deargen Inc., Daejeon, South Korea
77. Scriptomics, San Bruno, CA, USA
78. Department of Computer Science and Engineering, University at Buffalo, Buffalo, New York, USA
79. Data Science and Learning Division, Argonne National Laboratory, Lemont, IL, USA
80. Globus, The University of Chicago, Chicago, IL, USA
81. Turku Bioscience Centre, University of Turku, Turku, Finland
82. Research, Bioinformatics Division, Peter MacCallum Cancer Centre, Melbourne, Victoria, Australia
83. School of Medical Sciences, Orebro University, Orebro, Orebro, Sweden
84. Bioinformatics Division, Walter & Eliza Hall Institute of Medical Research, Melbourne, Victoria, Australia
85. School of Computing and Information Sciences, Florida International University, Miami, FL, USA
86. Biomolecular Sciences Institute, Florida International University, Miami, FL, USA
87. Sanford Burnham Prebys Medical Discovery Institute, Sanford Burnham Prebys Medical Discovery Institute, La Jolla, California, USA
88. Department of Biology, Brigham Young University, Provo, UT, USA
89. Department of Computational Biology, Indraprastha Institute of Information Technology, Delhi, Delhi, India
90. Department of Electrical Engineering, Korea Advanced Institute of Science and Technology, Daejeon, South Korea
91. McDermott Center for Human Growth and Development, University of Texas Southwestern Medical Center, Dallas, TX, USA
92. Salgomed, Inc., Salgomed, Inc., Del Mar, California, USA
93. Department of Biological Sciences, Graduate School of Science, The University of Tokyo, Bunkyo-ku, Tokyo, Japan
94. Department of Computer Science, KU Leuven, Leuven, Belgium
95. Translational Science and Experimental Medicine, Early Respiratory, Inflammation and Autoimmunity, R&D Biopharmaceuticals, AstraZeneca, Gaithersburg, MD, USA
96. Department of Plant Biotechnology and Bioinformatics, Ghent University, Ghent, Belgium
97. Department of Information Technology, IDLab, Ghent University, Ghent, Belgium
98. Department of Hematology/Oncology (Med V), Heidelberg University Hospital, Heidelberg, Germany
99. Department of Medical and Molecular Genetics, Indiana University School of Medicine, Indianapolis, IN, USA
