## Supplementary figures and images for "Multiple Myeloma DREAM Challenge Reveals Epigenetic Regulator *PHF19* As Marker of Aggressive Disease"

### Supplemental Figure 1

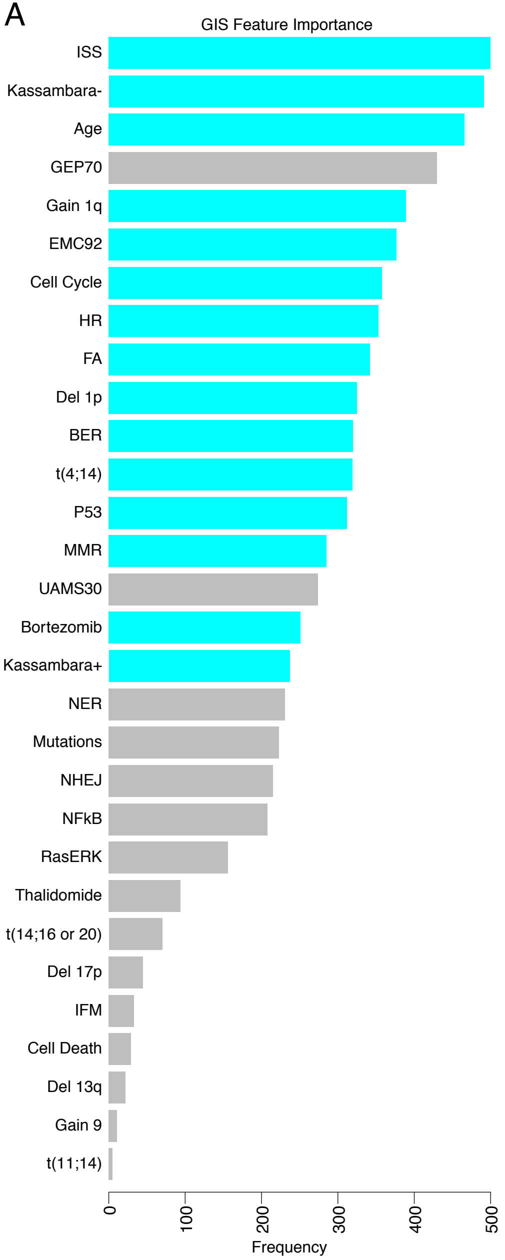

### Supplemental Figure 2

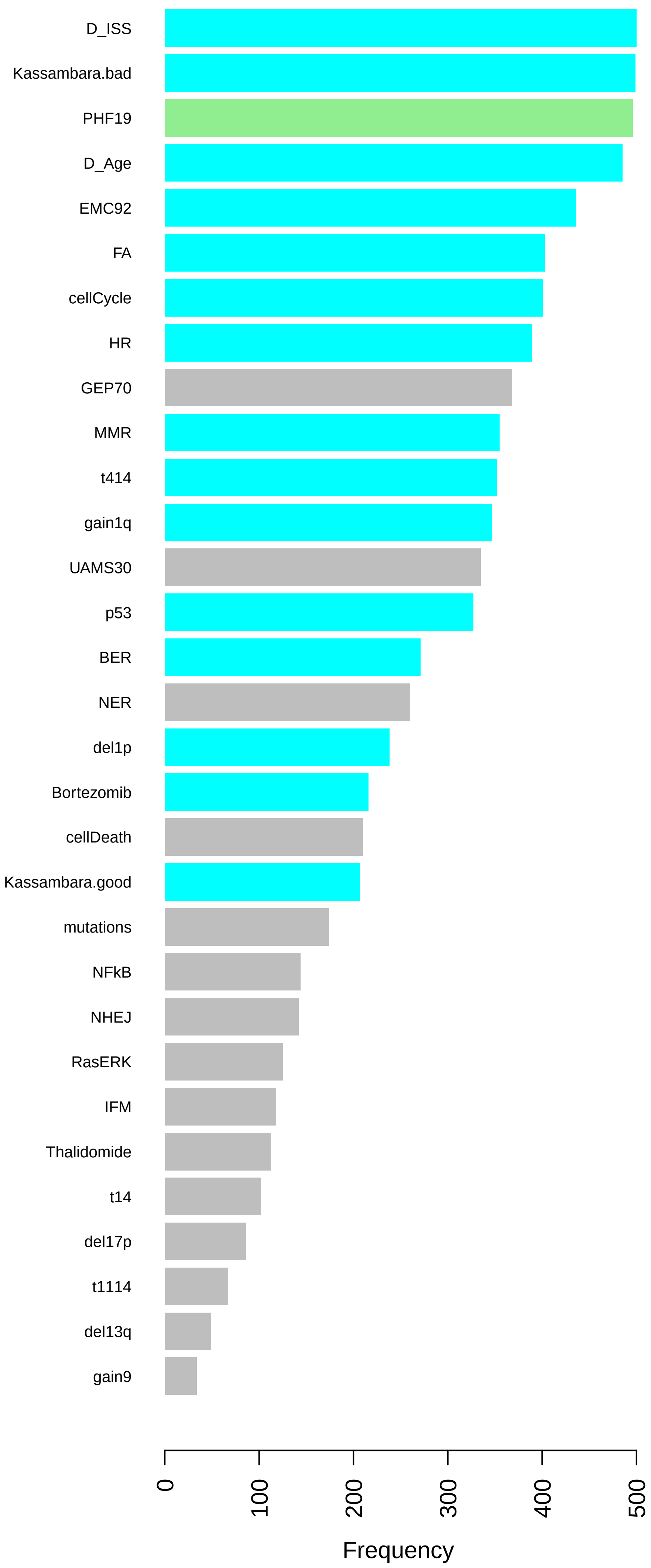

### Supplemental Figure 3

#### Cell Cycle

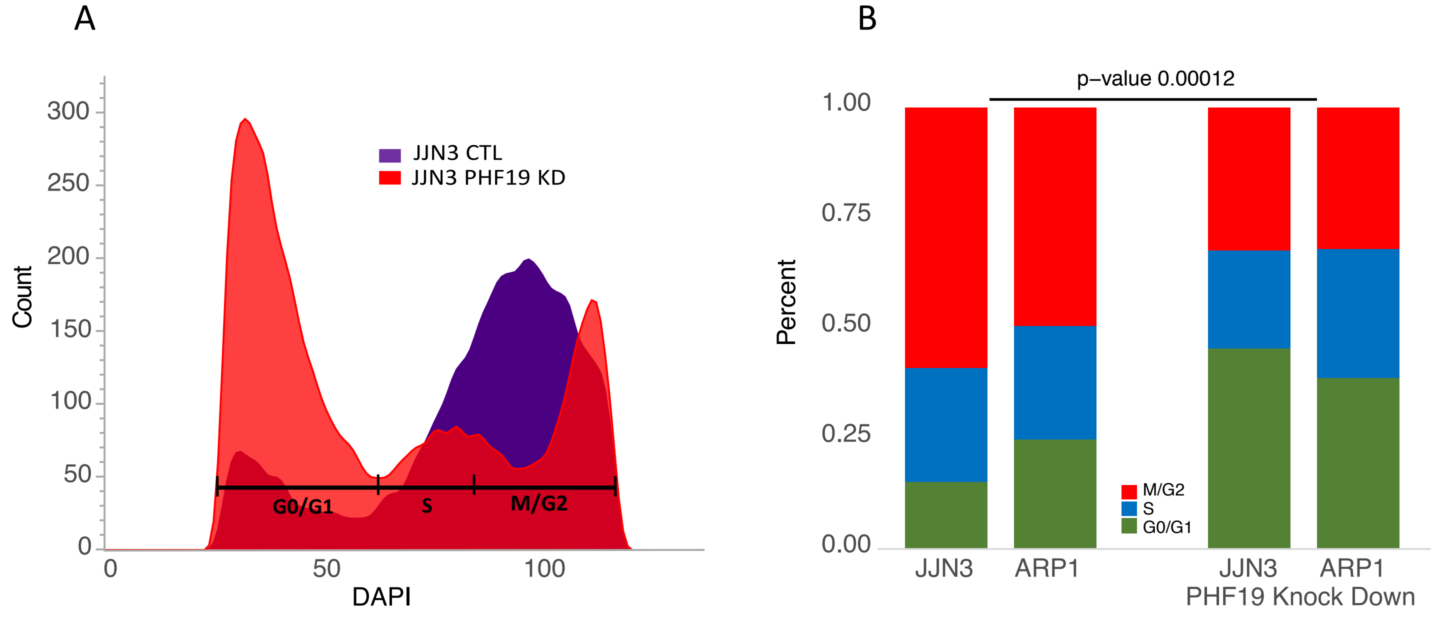


#### Apoptosis

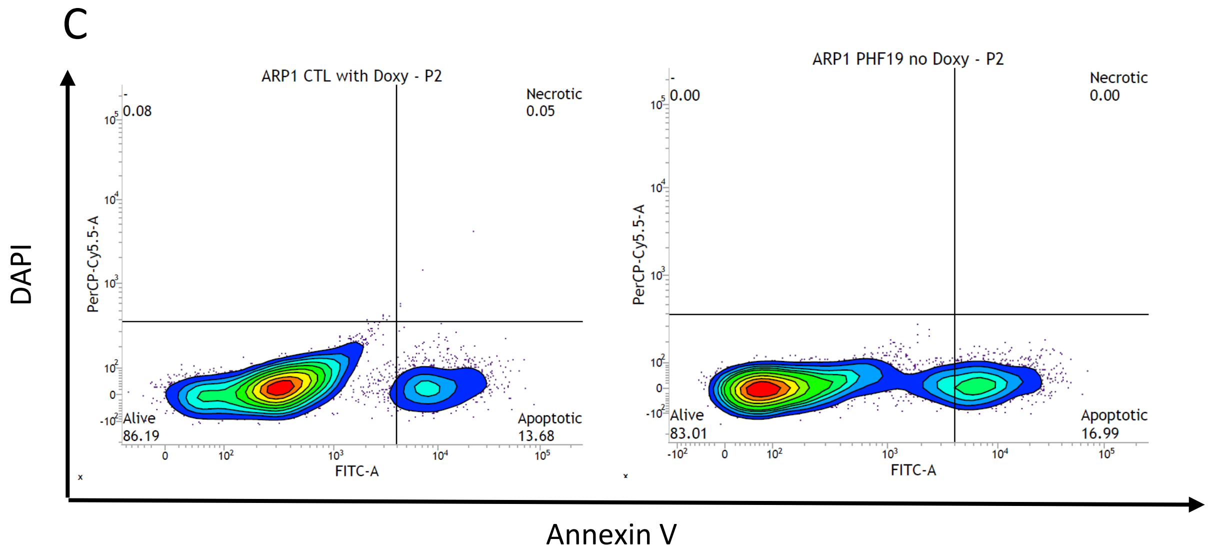


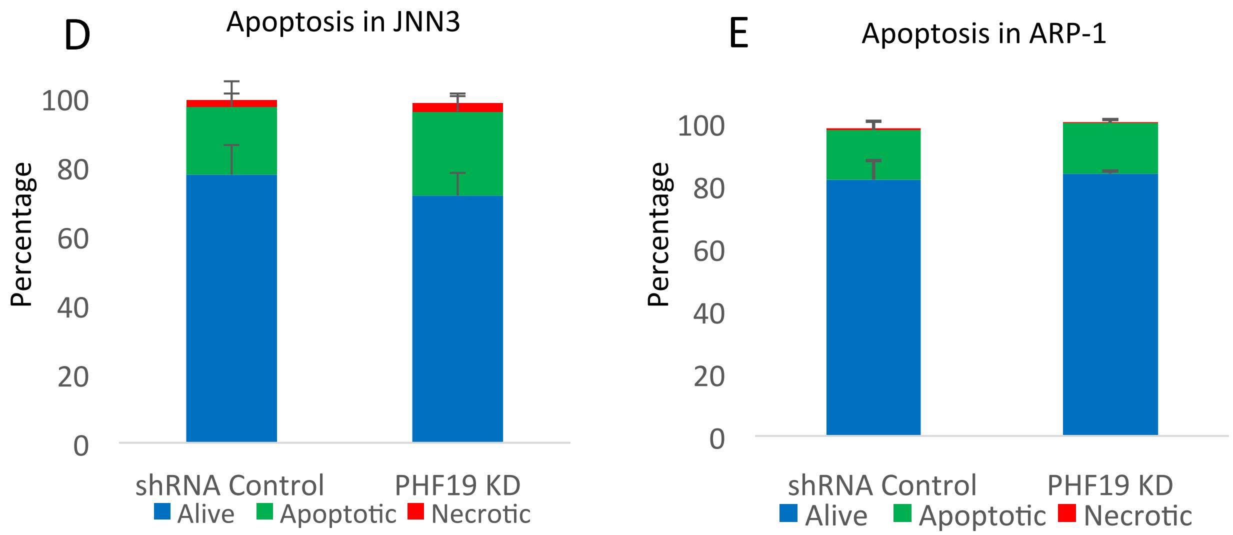


Supplemental Figure 3

### Supplemental Figure 4

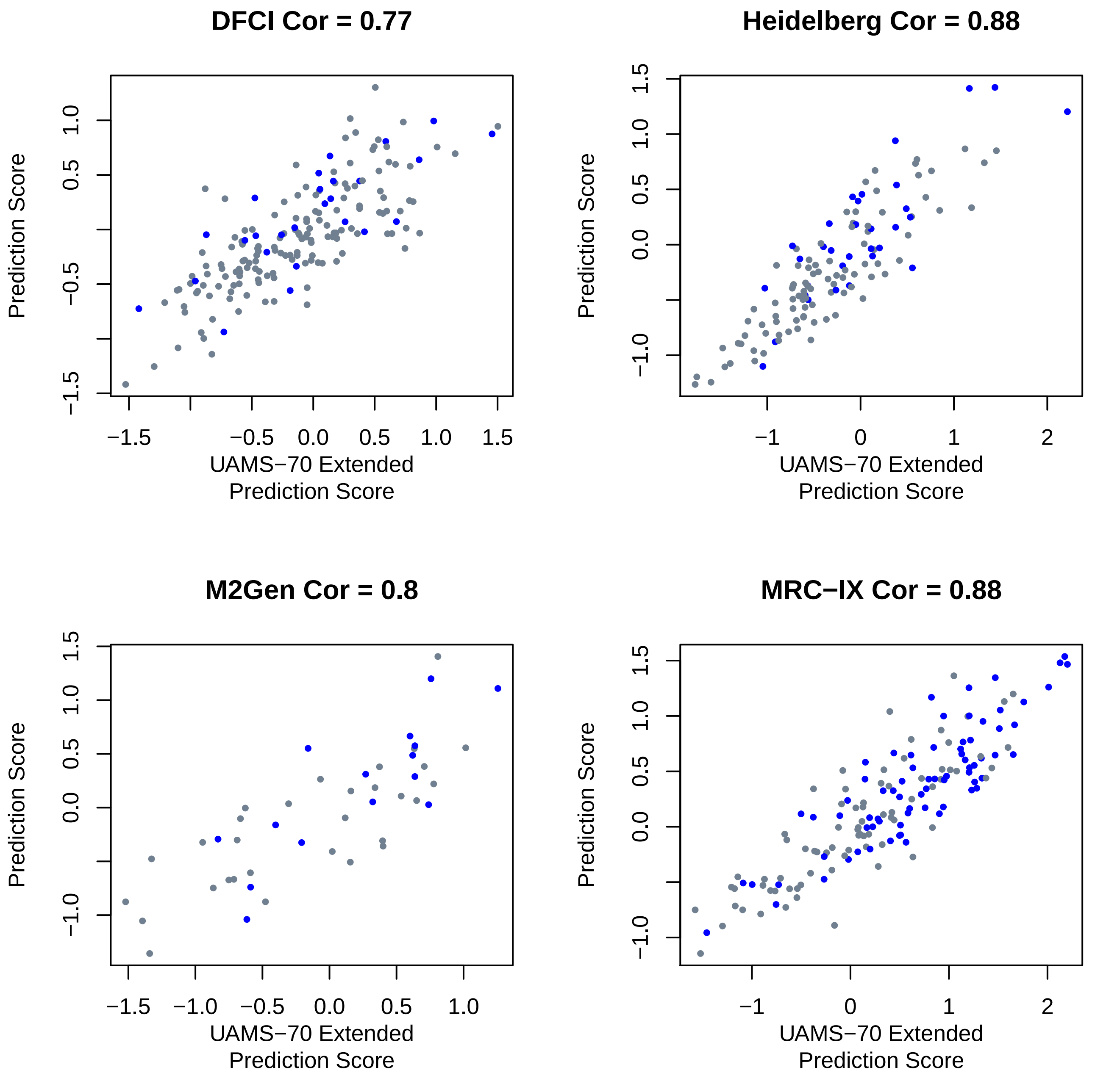
